# Task Related Neural Activity Following Primary Motor Cortical Ischemic Injury in Rats

**DOI:** 10.1101/2020.06.11.146266

**Authors:** Andrea R. Pack, Max D. Murphy, Scott Barbay, Randolph J. Nudo, David J. Guggenmos

## Abstract

Acquired injuries to primary motor cortex (M1) contribute to motor impairment and disability. Functional recovery is predicated on the reorganization of spared areas, which has been demonstrated through cortical motor map representations and neuroanatomical projection and termination patterns. The purpose of this study was to understand how neurophysiological outputs of spared motor areas relate to motor recovery of a skilled reach task following an ischemic infarct to M1. We examined changes in single unit activity within ipsilesional pre-motor (PM) and contralesional M1 cortices of rats during a behavioral task after a unilateral ischemic injury to ipsilesional M1. The data show a shift in neuronal firing patterns in the contralateral PM and ipsilateral M1 during behavioral recovery in lesion rats compared to a non-lesion control group, suggesting that spike-timing properties are altered in specific phases of the task, and that this altered activity may support spontaneous restoration of motor behavior.

**SIGNIFICANCE STATEMENT:** Following ischemic stroke to primary motor cortex (M1), motor recovery is associated with reorganization of spared cortical motor areas in injured and spared hemispheres. Currently, it is unclear how cortical plasticity within spared motor areas relates to motor recovery. This study examines how task-related neural activity within spared motor areas in rats correlates with motor restoration of a skilled reach task following an ischemic infarct to M1. The data suggest contralateral pre-motor and ipsilateral M1 alter their neural response profiles with respect to the timing of a motor task during recovery. To our knowledge, this is the first demonstration of a compensatory single-spike neurophysiological mechanism that may explain how remote, spared cortical areas contribute to functional recovery after M1 injury.

## INTRODUCTION

Ischemic stroke affects approximately 795,000 people in the United States every year and is one of the leading causes of disability (Benjamin et al., 2018). Research over the past two decades has shown that reorganization of the motor system, specifically within the spared pre-motor cortical areas that help with planning, initiating, and performing a behavior, occurs after a primary motor cortical lesion and may be the basis for the observed motor recovery (Dancause et al., 2005; Liu and Rouiller, 1999; Miyai et al., 1999; Rouiller et al., 1998). Despite the strong evidence for cortical plasticity driving motor recovery, it is unclear how activity within spared motor areas in both the injured and uninjured hemispheres relate to the restoration of motor deficits.

The rat motor cortex consists of two defined areas related to volitional motor control of the forelimb. The caudal forelimb area (CFA) is akin to primary motor cortex (M1) and has motor representations as shown via intracortical microstimulation (ICMS) (Donoghue and Wise, 1982; Neafsey et al., 1986). Rats can execute complex forelimb movements (Whishaw, 1996), resulting in reorganization of M1. The rostral forelimb area (RFA) functions somewhat analogous to that of pre-motor (PM) and supplementary motor areas (SMA) of primates, given its direct spinal projections and intracortical connectivity (Nudo and Masterton, 1988). Stimulation of PM can result in modulation of M1 outputs, indicating that PM has strong reciprocal cortico-cortical connections with M1, and its activity is an important component in controlling movement (Deffeyes et al., 2015; Rouiller et al., 1993). Recording studies examining the neurophysiological properties of PM neurons in rats have described similar firing patterns to M1 during a reach task, making it difficult to decipher how each area contributes to different aspects of the reach movement (Hyland, 1998; Saiki et al., 2014).

As previous cortical mapping studies have shown (Nishibe et al., 2010; Nudo et al., 1996), reorganization of spared motor areas following a lesion in M1 correlates with behavioral recovery. Furthermore, motor recovery coincides with a number of neurophysiological changes. Following skilled reach rehabilitative training, spared ipsilesional areas display changes in their motor representations. The significant increase in spared ipsilesional PM map size, seen in both humans and non-human animal models, correlates with behavioral recovery, suggesting that ipsilesional PM cortical areas take on at least part of the function of the damaged M1 to assist in recovery (Carey et al., 2002; Frost et al., 2003; Liepert et al., 1998; Nishibe et al., 2010). In addition to reorganization of ipsilesional motor areas, contralesional regions also undergo injury related changes (Marshall et al., 2000; Nhan et al., 2004). However, there is not a clear consensus whether the interhemispheric connections from contralesional M1 aid or hinder motor recovery.

Although there is strong evidence for motor cortical map reorganization and anatomical changes of spared motor areas related to behavioral recovery following stroke, there is little documentation of the neurophysiological response of spared motor areas during post-stroke recovery. We have employed chronic recording techniques to analyze single unit activity from these areas while the animal performed a skilled motor task. The purpose of this study was to understand how spared ipsilesional PM and contralesional M1 contribute to the restoration of reach and grasp following an ischemic infarct to the primary motor area. This was accomplished by recording extracellular activity of the spared PM contralateral to the preferred limb (corresponding to ipsilesional PM) and intact M1 ipsilateral to the preferred limb (corresponding to contralesional M1) during a reach task. This neural activity was examined after an ischemic stroke to M1 contralateral to the preferred (reaching) forelimb. The data show a shift in neuronal firing patterns in the contralateral PM and ipsilateral M1 during behavioral recovery in rats that sustained an ischemic stroke when compared to a non-lesioned control group, suggesting that these spared areas contributed to the restoration of reach and grasp behavior.

## METHODS

### Animals

Thirteen adult, male Long-Evans rats (weight = 312 – 353 grams, Harlan Laboratories, Inc., Indianapolis, IN) were acquired at three months of age. Rats were singly housed and provided food and water. The room was kept on a 12-h: 12-h light/dark cycle, and ambient temperature was maintained at 22 °C. Protocols for animal use were approved by the Kansas University Medical Center Institutional Animal Care and Use Committee and complied with the *Guide for the Care and Use of Laboratory Animals* (*Eighth Edition, The National Academies Press, 2011*). Rats (n=13) were assigned randomly to one of three groups: PM Control group (PM-C; n=4), PM Lesion group (PM-L; n=6), or M1 Control group (M1-C; n=3). The main experimental comparisons were between the PM-C and PM-L groups. Both groups had recording electrodes implanted in the rostral forelimb area (PM) contralateral to the reaching forelimb (Figure 1B; shown in green) and M1 ipsilateral to the reaching forelimb (Figure 1B; shown in orange). In addition, the PM-L group had an ischemic lesion in M1 contralateral to the reaching forelimb (Figure 1B; shown with six grey circles). The M1-C group was included in this study to confirm that the cortical activity recorded in contralateral M1 was consistent with reported behaviorally-related neural activity in previous studies (Hermer-Vazquez et al., 2004; Hyland, 1998). Thus, the M1-C group had recording electrodes implanted in the caudal forelimb area (M1) of both hemispheres (Figure 1B; shown in orange and blue).

**Figure 1.**
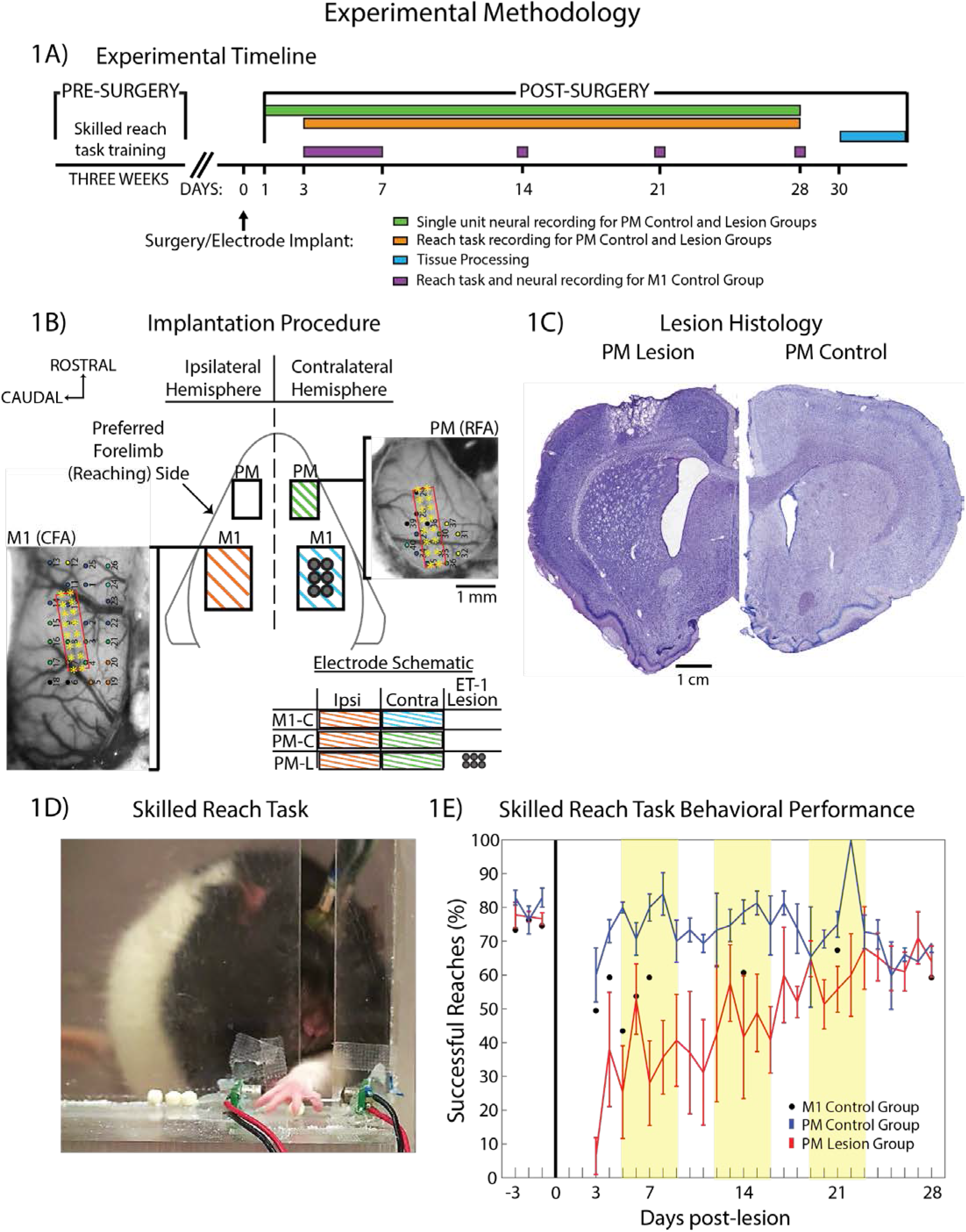
Experimental Design. 1A. Overview of experimental design. Rats were trained on a skilled reach task for up to four weeks. After reaching a 70% successful reach rate for 3 consecutive days, rats underwent surgery. PM Lesion (PM-L) and PM Control (PM-C) groups received microelectrode implants in PM contralateral and M1 ipsilateral to preferred forelimb and M1 Control (M1-C) group received bilateral M1 implants. Following electrode implants, rats were tested on the reach task on post-lesion days 3-28 for PM Control and PM Lesion groups (orange) and post-lesion days 3-7, 14, 21, and 28 for M1 Control group (purple) during a reach task session of 25 trials or a maximum of 20 minutes. 1B. Schematic of cortex showing the general topographic relationship between PM contralateral to preferred forelimb (contralateral PM (RFA); green), MI contralateral to preferred forelimb (contralateral M1 (CFA), blue), and M1 ipsilateral to preferred forelimb (ipsilateral M1 (CFA), orange) areas. Intracortical microstimulation (ICMS) was used to confirm location of the intact M1 and PM. M1-C group had one 16 site Neuronexus electrode array inserted into both ipsilateral (ipsi; orange) and contralateral (contra; blue) M1. PM-C and PM-L groups had one 16-shank microwire electrode array (Tucker Davis Technology - TDT) inserted into the contralateral PM (contra; green) and one into the ipsilateral M1 (ipsi; orange). The PM-L group had a focal, cortical ischemic infarct induced with endothelin-1 (ET-1 Lesion; six grey circles). The red outlined rectangle (array) and the 16 yellow asterisks (shanks) mark the placement of the implanted recording electrode. 1C. Coronal, cresyl violet stained brain slices of PM-L (left) and PM-C (right) groups at the level of M1. Staining reveals the ischemic lesion in M1 as a result of the ET-1 injections. 1D. Skilled Reach Task. Frame-by-frame video recordings were used for behavioral scoring and synchronization to neurophysiological data. 1E. Skilled reach task behavioral performance of M1-C (black), PM-C (blue), and PM-L (red) groups. Error bars at each time point represent the standard error of the mean. Days −3, −2, and −1 represent the mean baseline performance across the three days prior to electrode implantation (M1-C group = 74.63 +/- 1.68% (SEM), PM-C group = 80.69 +/- 3.04 (SEM), and PM-L group = 77.22 +/- 2.37% (SEM)). The yellow blocks represent the three time points analyzed in the behavioral results: Day 7 +/- 2 = Week 1, Day 14 +/- 2 = Week 2, Day 21 +/- 2 = Week 3.

### Behavioral training

Rats were trained on a skilled reach task (Whishaw et al., 1991) for up to four weeks before surgery (Figure 1D). Each rat was placed within a 30-cm x 30-cm x 53-cm Plexiglas box with a 1-cm-wide slot allowing access to a tray where food pellets were placed. Forelimb preference was determined at the beginning of the skilled reach task training (Hsu and Jones, 2006). Rats were trained to reach out of the slot, grasp a food pellet (45 mg; Bio-Serv. Flemington, NJ) located 2 cm away on a horizontal shelf, and bring the food pellet to the mouth (successful retrieval). Trials were recorded with a digital camcorder for playback and analysis. Rats were required to obtain at least a 70% successful reach rate (60 trials per day) for three consecutive days (baseline behavior) before inclusion into the study. Throughout the duration of the study, food intake was monitored on a scheduled feeding regimen to encourage performance on the skilled reach task (Bury and Jones, 2002).

### Surgery

Each rat was anesthetized initially with isoflurane (to effect) followed by ketamine (80-100 mg/kg, intraperitoneal (IP)) and xylazine (5-10 mg/kg, intramuscular (IM)). Anesthesia was maintained throughout the procedure with repeated bolus injections of ketamine (10-15 mg/kg/ IM) as needed. After the rat was secured in the stereotaxic frame, bupivacaine (0.5-1.0 mL) was delivered subcutaneously (SC) to the scalp/incision area and penicillin (0.15 mL; G benzathine and G procaine, 300 K/mL) was delivered SC under the skin of the back. A rectal probe attached to a homothermic blanket (Harvard Apparatus, Holliston, MA) was used to maintain core body temperature at approximately 36.7°C throughout the experiment. Ocular ointment (Paralube Vet Ointment, ParmaDerm, Princeton, NJ) was then applied to the eyes. Using aseptic techniques, the scalp was incised along the midline of the skull. The temporalis muscle was retracted and a craniectomy was made based on stereotaxic coordinates relative to bregma over the PM contralateral to the reaching forelimb (anterior-posterior (AP) +3.5 mm, medial-lateral (ML) +2.5 mm) and the M1 ipsilateral to the reaching forelimb (AP +0.5 mm, ML + 3.5 mm) (Kleim et al., 1998; Nishibe et al., 2010) (Figure 1B). To control brain edema after the craniectomy, the muscles of the neck overlying the cisterna magna were reflected and a small puncture was made in the spinal dura (foramen magnum) to allow for CSF drainage. Six holes (three on either side of the skull midline) were drilled caudal to the M1 craniectomies for placement of stainless steel screws used during the procedure to ground the electrodes and anchor them to the skull with dental acrylic. The dura was then removed over the exposed PM and M1. ICMS was used to identify the forelimb motor representations (Figure 1B). Following completion of the surgical procedure, the incision was closed with sterile 4.0 silk surgical suture and treated topically with EMLA cream (lidocaine and prilocaine cream, Dechra Veterinary Products, Overland Park, KS), and triple antibiotic ointment. The rat was removed from the stereotaxic device and the temperature probe removed and placed within the homeothermic blanket, upon which the animal was monitored for anesthetic recovery. Each rat received an additional injection of penicillin 0.15 mL SC (G benzathine and G procaine 45,000 IU), and then was moved to a recovery area until the rat was able to maintain an upright posture and locomote freely. Once stabilized, the rat was returned to its home cage. Post-surgery analgesics were given to the rat during the 72 hours after surgery [four SC injections of Buprenorhpine (0.05-0.10 mg/kg every 12 hours; four doses of Acetaminophen (20-40 mg/kg, oral) every 12 hours].

### ICMS motor mapping

ICMS procedures were used to identify the primary motor and premotor cortex forelimb fields as described previously (Kleim et al., 2002; Kleim et al., 1998; Nudo et al., 1996). A digital microphotograph of the cortical surface was acquired and transferred to a graphical illustration program (CANVAS; Canvas GFX, Inc., Fort Lauderdale, FL) and overlaid with a 250 μm grid. Responses at the grid intersections (avoiding surface vasculature) were recorded and used to inform microelectrode array placement (Figure 1B).

### ET-1 lesion

After identification of the motor forelimb fields in the cerebral cortex, rats assigned to the PM-L group had six 0.7-mm diameter holes drilled over the M1 contralateral to the reaching forelimb (AP +1.5, +0.5, −0.5 mm and ML +2.5, +3.5 mm). A focal, cortical ischemic infarct was induced with the vasoconstrictor, endothelin-1 (ET-1; 0.3 μg ET-1 dissolved in 1 μl sterile saline (~120 pmol); Bachem Americas, USA) (Adkins et al., 2004) (Figure 1B, shown in gray). Each hole was injected with 0.33 μL of ET-1 through a glass micropipette (160 μm o.d.) attached to a Hamilton syringe with the tip located at 1.5 mm below the cortical surface. Injections were made at a rate of 4nl/sec via an electronic microsyringe injector (UltraMicro Pump III, World Precision Instruments, Sarasota, FL). The injection pipette remained in place for an additional 5 minutes after injection to minimize backflow of ET-1 during pipette removal from the cortex (Barbay et al., 2013).

### Microelectrode array implant

Following ICMS and ET-1 lesion procedures, skull screws (Pan Head Phillips head machine Screws, 0-80 thread, 1/8” length, stainless steel, McMaster-Carr, Elmhurst, IL) were placed in the six screw holes to ground the electrodes and to act as an anchor for dental acrylic applied to the skull. For the PM-C and PM-L groups, two chronic, recording Tucker-Davis Technologies (TDT; Alachua, FL) microelectrode arrays (16 shank, 2×8 electrode, 50 μm wire diameter, 45° tip angle, 250 μm electrode spacing, 500 μm row spacing) were implanted in the PM contralateral to the reaching forelimb and M1 ipsilateral to the reaching forelimb. In the M1-C group, two chronic, recording microelectrode arrays (NeuroNexus Technologies, Ann Arbor, MI; A4×4-3mm-100-125-703) were implanted bilaterally in M1. On-line recording of activity from the microelectrode arrays was used to determine a depth with high neuronal firing on the majority of channels as the array was lowered into place with a micropositioner (at a depth ~1550 μm perpendicular to the cortical surface). A quick-curing polymer was used (Kwik-Cast, WPI, Sarasota, FL) to cover the cortex, fix the arrays in place, and seal the cranial opening. Dental acrylic was then used to add additional anchoring to the skull by adhering to the stainless steel screws placed in the skull. The acrylic also served as an additional protective barrier over the polymer.

### Behavioral procedures and neurophysiological data collection

Each rat was tethered to the implanted arrays through a motorized commutator that allowed free movement within the testing chamber. Neuronal activity was recorded using TDT hardware and a custom TDT OpenEx system circuit with a bandpass filter between 300 and 5,000 Hz and sampling rate at 25 kHz. Following the lesion and array implants, each rat was tested on the reach task on post-lesion days 3-28 for PM-C and PM-L groups and post-lesion days 3-7, 14, 21, and 28 for M1-C group during a reach task session of 25 trials or a maximum of 20 minutes (Figure 1A). When the rat reached through the 1 cm Plexiglas slot, a photobeam was broken (beam break). A user-operated button was pressed each time a rat had a successful reach to differentiate successful and unsuccessful reaches. The TDT circuit collected raw waveforms of the extracellularly-recorded neuronal spikes on each channel for each electrode site as well as time stamps for the beam-break and button press (successful reaches). All data were processed offline using custom MATLAB (Mathworks, Natick, MA) scripts to determine the correlation between phases of the behavior and neuronal activity. A repeated-measures mixed-model ANOVA was used to determine the effects of group, time and group-by-time interactions on behavioral performance scores (JMP, Care, NC). Reaching performance scores (percentages) were transformed using an arcsine transformation to adjust for normality.

### Histology

Rats were perfused transcradially with normal saline followed by 3% paraformaldehyde. Brains were post-fixed in 30% glycerol solution. Coronal sections were cut at 50 μm and stained with cresyl violet. Sections were mounted to slides, dehydrated, and cover slipped (Figure 1C).

### Behavioral Analysis - video scoring and alignment

Videos for documentation and offline analysis of behavioral performance and alignment to neurophysiological data were recorded at 30 frames per second. All offline scoring and post-processing steps were conducted using custom MATLAB functions. Videos were manually scored to obtain the frame of reach and grasp onset. Reach onset was defined as the first frame in which the rat could be seen moving its forelimb out of the box. Grasp onset was defined as the first frame in which the rat could be seen closing its forepaw (Figure 1D). Trials were only considered where both a reach and a grasp could be identified (thus, trials where the rat reached but did not grasp because a pellet was not present were not considered as an unsuccessful trial). Trials were considered successful if the rat grasped the pellet and brought it back to its mouth without dropping it.

To align the video frame time series and neural data time series, beam breaks that were sampled synchronously with the neural data were manually aligned on a per frame basis such that the occurrence of beam breaks consistently corresponded visually to times when the rat was reaching across the plane of the beam. Recordings where alignment was not readily apparent were discarded.

### Neurophysiological data reduction and processing

Based on the quality and availability of neurophysiological recordings, 2-3 days per animal surrounding the following three time points were considered for analysis: Day 7+/- 2 (‘Week 1’), Day 14+/- 2 (‘Week 2’), and Day 21+/- 2 (‘Week 3’). For each recording, snippets of multi-unit activity corresponding to a range from −500 ms before to 150 ms relative to each reach onset were examined (Figures 2A,B). Trial multi-unit recordings were then individually inspected visually for the presence of chewing or whisking artifact, which were clearly observed as periodic bursts of activity within the filtered multi-unit signal. Trials containing such artifact, as well as individual microwire channels that appeared to be broken based on their signal profile were excluded from further analysis. After artifact exclusion, the remaining channels were virtually re-referenced to overcome common noise sources (Ludwig et al., 2009). Following re-referencing, trials for each channel were visually inspected to ensure that re-referencing did not introduce artifact. Spikes were detected offline (Maccione et al., 2009) and considered in 1 ms bins for the remainder of the analyses. After detection, spikes were automatically sorted using superparamagnetic clustering (Quiroga et al., 2004). The resulting clusters were manually combined or excluded based on the visually assessed quality of average unit spike profiles and variability. Single-units were identified within the course of a single recording, but were not matched over multiple consecutive recordings. A total of 264 single units were identified over 68 recording sessions (Table 1).

**Figure 2.** Data Extraction and Analysis. 2A,B. Raster plots for successful grasp trials (2A) and unsuccessful grasp trials (2B). The plots show neuronal spike activity beginning 500 ms prior to the onset of paw closing (grasp onset; vertical line) to 150 ms after the grasp onset. Each row represents a different grasp trial. Each tick mark corresponds to an action potential (spike) of a single semi-automatically characterized neural unit. Spikes detected in the bandpass filtered (300 Hz to 5000 Hz) extracellular signal were clustered into neural units using a combination of automatic and manual scoring to reject clustered artifacts. 2C,D. Rate estimation for successful grasp trials (2C) and unsuccessful grasp trials (2D). To capture fluctuations in the spike activity on time scales that may be faster or slower than the width of a given histogram period, we used an adaptive smoothing algorithm to estimate continuous fluctuations in trial-averaged neural activity. The Gaussian kernel changes bandwidth depending on the timescale of changes in the neural activity. 2E,F. Individual unit spike rate trajectories for successful grasp trials (2E) and unsuccessful grasp trials (2F). The bold lines in E and F (blue and red, respectively) represent the trial-averaged spike rate for the unit whose rasters are displayed in A and B, respectively. In order to estimate how disperse the excited units were in the time of peak onset relative to grasp alignment, the inter-quartile range (IQR; grey box) was computed from the set of maxima onset times obtained from each included unit. The peak of the bold line represents the upper bound on the IQR for the time of absolute rate maximum of each unit. The thin line represents the unit whose rate maximum occurs at the lower bound of the IQR for rate maximum onset. In order to study units that were excited relative to the behavior, only units with average firing rates of at least 5 spikes per second and a significantly (α = 0.005) higher maximum rate during grasp onset in comparison to maximum rate during random event alignment were included for further analysis. Units not meeting this criterion (shown in dashed line) were excluded. 2G, H. Spike rate ensemble averages for successful grasp trials (2G) and unsuccessful grasp trials (2H). The average of each included individual unit trajectory (i.e. blue and red, bold and thin solid lines in 2E,F) was used to estimate the average spiking excitation relative to grasp alignment. The thick blue and red lines represent the average spike rate of units during alignment to successful and unsuccessful grasps, respectively. Dot-dashed lines represent the standard error of the mean (SEM). The bold black line represents the resampled average rate trajectory for all units with randomized grasp alignment times. Black dash-dot lines represent SEM for the same trajectories. Note that the SEM of the randomized trials is smaller because all recorded units that fired with a rate of at least 5 spikes per second were included in the resampling procedure. In order to assess the duration of time for which the ensemble maintained an average excited state relative to the grasp alignment, periods in which the ensemble average was greater than the average resampled ensemble average plus 10 times its SEM were computed (light blue and red boxes). Table 1 lists counts of included units, IQR values, and the time period where neural units had spike rates >10 times SEM of randomly aligned data (>10 SEM) for Figures 3-5.

**Table 1:** Summary of identified neural units, inter-quartile ranges (IQR), and the time period where neural units had spike rates >10 times SEM of randomly aligned data (>10 SEM) for each analysis completed in Figures 3-5. IQR is the time range within which included units (neural unit count) were most active during skilled the skilled reach task (500 ms prior and 150 ms after the grasp onset). Negative values (-) used to define IQR and >10 SEM start and end times represent any time point that occurred prior to grasp onset, whereas positive values (+) used to define the IQR and >10 SEM start and end times represent any time points that occurred after grasp onset.

|  | Contralateral |  |  |  |  |  |  |  |
| --- | --- | --- | --- | --- | --- | --- | --- | --- |
|  | Neural Unit Count |  | IQR (ms) |  |  | >10 SEM (ms) |  |  |
|  | Aligned | Random | Start Time | End Time | Total Time | Start Time | End Time | Total Time |
| Successful | 5 | 22 | -168 | -52 | 116 | -69 | -40 | 29 |
| Unsuccessful | 12 | 20 | -186 | -61 | 125 | N/A | N/A | 0 |
|  | Ipsilateral |  |  |  |  |  |  |  |
|  | Neural Unit Count |  | IQR (ms) |  |  | >10 SEM (ms) |  |  |
|  | Aligned | Random | Start Time | End Time | Total Time | Start Time | End Time | Total Time |
| Successful | 6 | 45 | -163 | +130 | 293 | +25 | +120 | 95 |
| Unsuccessful | 15 | 48 | -324 | -3 | 321 | N/A | N/A | 0 |

**Contralateral PM Activity (Figure 4)**
|  | PM-C |  |  |  |  |  |  |  |
| --- | --- | --- | --- | --- | --- | --- | --- | --- |
|  | Neural Unit Count |  | IQR (ms) |  |  | >10 SEM (ms) |  |  |
|  | Aligned | Random | Start Time | End Time | Total Time | Start Time | End Time | Total Time |
| Successful | 15 | 51 | -466 | +65 | 531 | N/A | N/A | 0 |
| Unsuccessful | 22 | 46 | -382 | +67 | 449 | N/A | N/A | 0 |
|  | PM-L |  |  |  |  |  |  |  |
|  | Neural Unit Count |  | IQR (ms) |  |  | >10 SEM (ms) |  |  |
|  | Aligned | Random | Start Time | End Time | Total Time | Start Time | End Time | Total Time |
| Successful | 40 | 212 | -157 | +115 | 272 | -35 | +120 | 155 |
| Unsuccessful | 106 | 194 | -396 | +3 | 399 | N/A | N/A | 0 |

**Ipsilateral PM Activity (Figure 5)**
|  | PM-C |  |  |  |  |  |  |  |
| --- | --- | --- | --- | --- | --- | --- | --- | --- |
|  | Neural Unit Count |  | IQR (ms) |  |  | >10 SEM (ms) |  |  |
|  | Aligned | Random | Start Time | End Time | Total Time | Start Time | End Time | Total Time |
| Successful | 54 | 120 | -23 | +124 | 147 | +24 | +120 | 96 |
| Unsuccessful | 57 | 90 | -350 | 102 | 452 | N/A | N/A | 0 |
|  | PM-L |  |  |  |  |  |  |  |
|  | Neural Unit Count |  | IQR (ms) |  |  | >10 SEM (ms) |  |  |
|  | Aligned | Random | Start Time | End Time | Total Time | Start Time | End Time | Total Time |
| Successful | 51 | 256 | -150 | +84 | 234 | -122 | +120 | 242 |
| Unsuccessful | 107 | 248 | -322 | +88 | 410 | N/A | N/A | 0 |

### Rate estimation and identification of task-related neurons

To gauge the activity of individual neurons for a given alignment condition (i.e., successful or unsuccessful grasps and reaches), smoothed continuous estimates of the spiking rate profile for each neuron were obtained. Each trial within an alignment condition was considered to be independent; thus, spike times relative to the trial onset could be aggregated into a single trial wherein individual spikes were assumed to be independent of one another. Using this assumption, the firing rate of an individual neuron was estimated using an adaptive kernel density smoother (Figures 2C,D) (Shimazaki and Shinomoto, 2010a). This method has an adaptive stiffness parameter that affects how rapidly the Gaussian smoothing kernel is updated. We found that the stiffness constant tended to skew to the left, with a mean of 0.86 and median of 0.77. The resulting estimate of the number of spikes per second, per trial, was then segmented into 1 ms time bin increments for the specified trial duration, resulting in discrete rate estimate functions with 651 samples (Figures 2E,F; bold line). Firing rate profiles of individual neurons were then included for further analysis if the average firing rate was at least 5 spikes per second, and if there were at least 5 trials for a given alignment condition on that recording. Only units demonstrating a peak in spike rate greater than a pre-determined threshold relative to alignment were included for subsequent analysis. The threshold was set on a per-unit basis, depending on the average number of spikes observed per trial, in order to account for peaks in units with lower activity and to take into account the effect of the rate estimator. For each unit, 1000 surrogate trials were generated using the number of spikes observed relative to each alignment. Surrogate trials assumed a uniform random distribution of spike times. After applying the rate estimator to each surrogate, the maximum rate was obtained and the maximum rate for the observed unit was compared to the 95^th^ percentile surrogate maximum rate (Saiki et al., 2014). Units with a lower maximum rate were excluded (Figures 2E,F; dashed line). It should be noted that this identification strategy tends to incorporate units that are excited relative to behavioral alignment, rather than those which are inhibited. Rate estimates for units from the same cortical area and treatment condition were then averaged together (Figures 2G,H; bold line) yielding the ensemble average neural rate trajectory for each condition (Figures 3-5). The standard error of the mean (SEM, Figures 2G,H; dashed lines) was also estimated in order to measure the relative dispersion in firing rates of all included units within an ensemble throughout the time course of the grasp.

**Figure 3:** Spike rate ensemble averages during reach task execution - MI Control (M1-C) group. A, B: Contralateral M1 spike rates in successful (A) and unsuccessful (B) trials. C, D: Ipsilateral M1 spike rates in successful (C) and unsuccessful (D) trials. The dark blue (A, C) and red (B, D) lines represent the average spike rates during successful and unsuccessful trials, respectively. Black lines represent the average spike rate when alignment times were randomly aligned. Dot-dashed lines represent the standard error of the mean (SEM). Light-blue vertical bars (A, C) represent time periods when the average spike rate exceeded the randomly aligned rate by more than 10 times SEM. Vertical dashed black line at time zero depicts time of grasp onset. Spike rates met the 10 SEM criterion in contralateral M1 prior to the successful grasps, and in ipsilateral M1 after successful grasps. Spike rates did not meet the criterion at any time in either hemisphere during unsuccessful trials.

**Figure 4:**
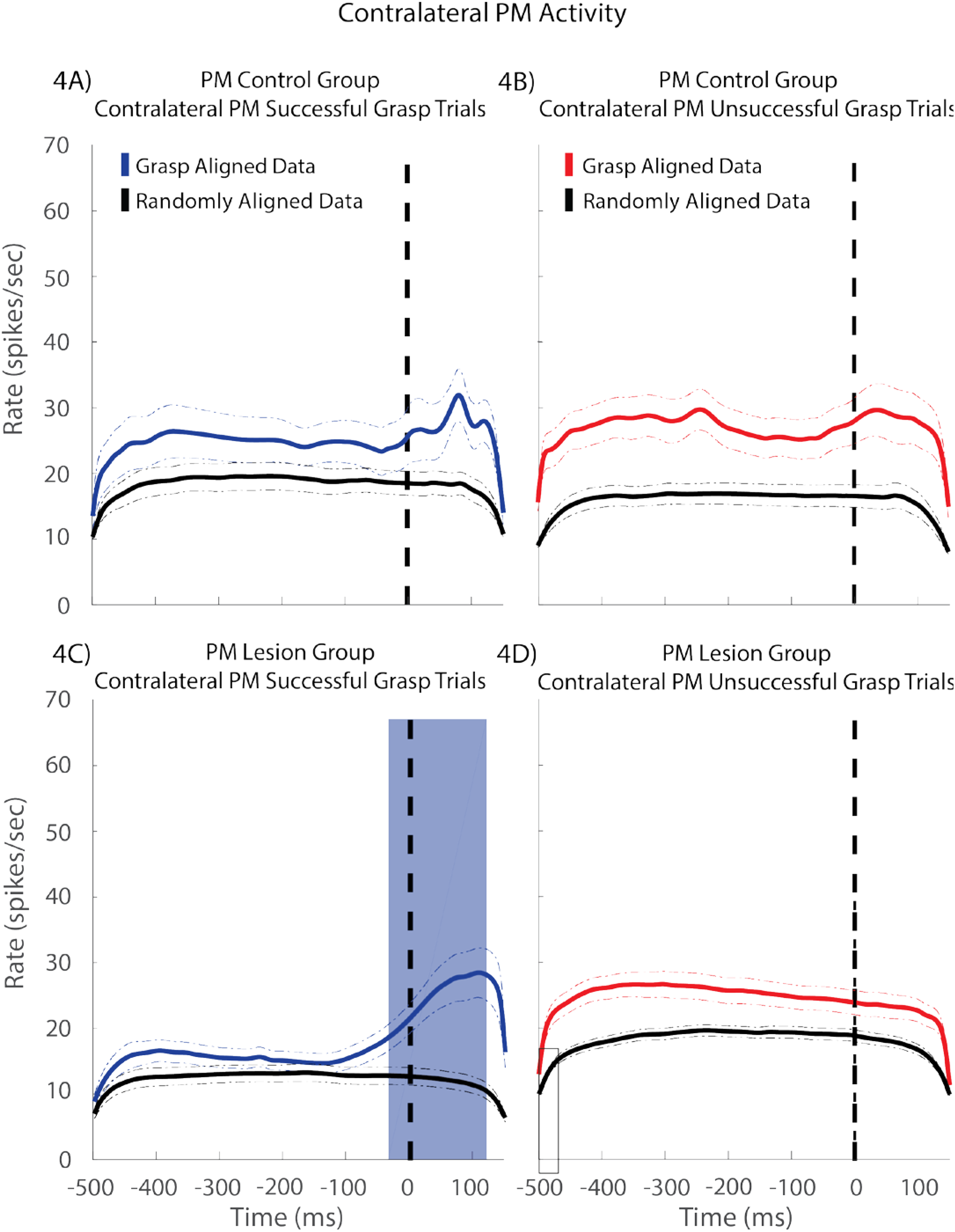
PM Control (PM-C) and PM Lesion (PM-L) groups – contralateral to reaching forelimb PM activity. Neural excitation demonstrated by spiking of single-and multi-units within contralateral (ipsilesional to the preferred forelimb) M1 of PM-C and PM-L animals was split into successful (left; Figures 4A,C) and unsuccessful (right; Figures 4B,D) grasp trials. The thick blue and red lines represent the average spike rate of units during alignment to successful and unsuccessful grasps, respectively. Black lines represent the average spike rate when alignment times are randomly shuffled. Dot-dashed lines represent the standard error of the mean (SEM). The light-blue (Figure 4C) box represents the region where the average rate of all units is greater than the average re-shuffled rate plus 10 SEM. Vertical dashed black line at time zero depicts time of grasp onset. Unsuccessful trials in both PM-C and PM-L rats showed similar profiles, with a general increase in activity across the entire time range relative to random event alignment. PM-L successful trials showed an increase in activity starting approximately 50 ms prior to the grasp onset and persisting at least 150 ms following grasp onset compared to PM-C successful trials, which show a smaller increase of activity after the grasp onset.

**Figure 5:**
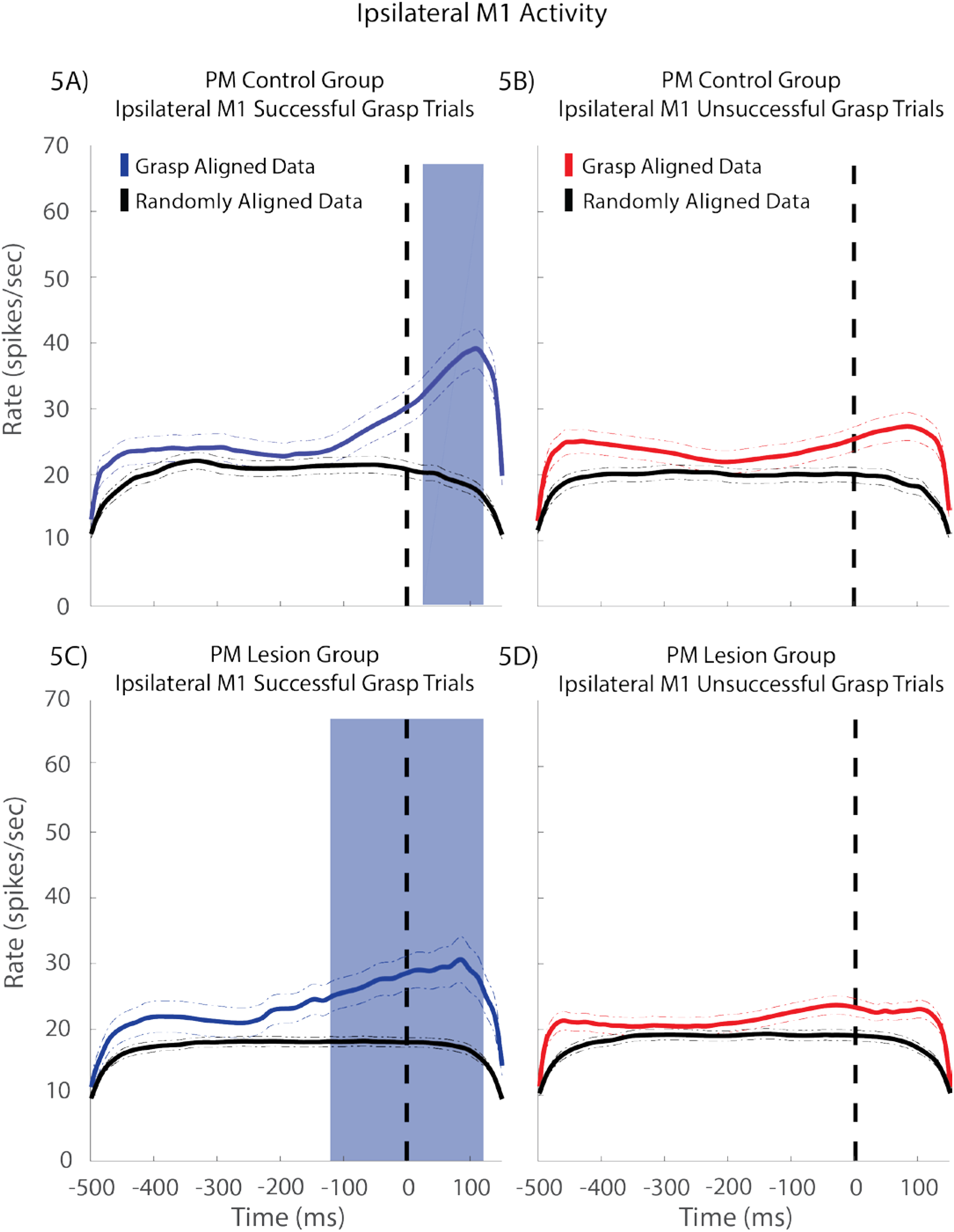
PM Control (PM-C) and PM Lesion (PM-L) groups – ipsilateral to reaching forelimb M1 Activity. Neural excitation demonstrated by spiking of single-and multi-units within ipsilateral (contralesional to the preferred forelimb) M1 of PM-C and PM-L animals was split into successful (left; Figures 5A,C) and unsuccessful (right; Figures 5B,D) grasp trials. The thick blue and red lines represent the average spike rate of units during alignment to successful and unsuccessful grasps, respectively. Black lines represent the average spike rate when alignment times are randomly shuffled. Dot-dashed lines represent the standard error of the mean (SEM). Light-blue (Figures 5A,C) boxes represent regions where the average rate of all units is greater than the average re-shuffled rate plus 10 SEM. Vertical dashed black line at time zero depicts time of grasp onset. There is a similar pattern of activity between PM-C and PM-L groups during successful reaches (Figures 5A,C, respectively), with slight differences in the expected time at which activity increases prior to grasp onset and the time peak activation occurs after grasping the pellet. Unsuccessful trials in both PM-C and PM-L (Figures 5B,D, respectively) rats showed similar profiles; The ensemble rate average (thick blue and red lines) never fired 10 SEM above the resampled average (thick black lines).

### Significance estimation of rate profiles

To gauge the significance of rate profiles, alignment times were randomly re-shuffled and trials were re-sampled from the recorded data. The re-shuffling procedure obliterates spike rate peaks due to task-relatedness while preserving any inherent bias in rate profile shape due to the inherent inter-spike interval of the recorded spike profiles (Grün and Rotter, 2010). After re-shuffling, randomly aligned surrogate trials were processed using the same rate estimation and exclusion steps as the raw data, to account for biases introduced during processing. In subsequent criteria, “rate profiles” refer to the average (mean) spike rate from all trials recorded from a single session, for an individual unit. Ensemble averages therefore refer to all rate profiles fitting a given condition (combinations of lesion vs. control, RFA vs. CFA, and/or week number). For example, an ensemble could consist of all units belonging to animals from the PM-L group. To identify the dispersion of periods in which individual units included in an ensemble were most active, the ensemble interquartile range (IQR) was computed from the onset times of each rate profile maximum relative to grasp alignment. In addition, because the IQR does not contain information about peak magnitude, we also defined time periods when ensemble average spike rate fluctuations indicated significant excitation. This was defined as the time period in which the grasp-aligned ensemble mean spike rate was greater than the random-alignment ensemble mean by at least 10 times the SEM of the random-alignment ensemble (Grün and Rotter, 2010; Hyland, 1998). The conservatively high threshold reflects the focus on identifying periods around behavior in which a majority of units demonstrate relatively focal excitation. It is higher than other threshold criteria (i.e. Hyland, 1998) because there is no additional specification to exceed threshold for any number of consecutive samples. It should be noted that both methods are focused on identification of significantly excited units, and therefore significantly inhibited units are not considered in the scope of this study.

## RESULTS

### M1 ischemic injury impairs skilled reach task performance

A skilled reach task was used to assess behavioral performance following an ischemic infarct to M1 contralateral to the trained forelimb. Prior to the lesion, the rats were trained to criterion baseline performance (70%). The mean baseline success rate was 80.69 +/- 9.75% (SEM) and 77.22 +/- 7.96% (SEM) for the PM-C and PM-L groups, respectively. Since rats were tested on slightly different post-infarct days, data were grouped into the following time periods: Baseline, Week 1 (Day 7 +/- 2), Week 2 (Day 14 +/- 2) and Week 3 (Day 21 +/- 2). A repeated measures mixed-model ANOVA revealed a significant effect of Group [F(1,8) = 5.8867; p = 0.0414] and Time [F(3,24) = 3.6178; p = 0.0276], but no Group by Time interaction [F(3,24) = 2.0601; p = 0.1322] on behavioral performance (Figure 1E), demonstrating that the infarct had a significant effect on behavioral performance in the PM-L group relative to the PM-C group. Since the M1-C group was included in this study to verify typical timing of behaviorally-related spike firing, and behavioral performance data from these rats were collected less frequently, it was not included in the statistical comparison of behavioral performance.

### Temporal relationship of spike timing to execution of motor task

As described in the Methods, the firing rates of individual neurons relative to the grasp onset were estimated using an adaptive kernel density smoother (Shimazaki and Shinomoto, 2010b). The solid lines defined as ‘Grasp Aligned Data’ in Figures 3-5 depict the time course of an ensemble average spike rate for identified units relative to the grasp onset, whereas the solid black lines defined as ‘Randomly Aligned Data’ depict the time course of an ensemble average spike rate using random time points for alignment. Data were analyzed separately for successful trials (trials in which the rat grasped and brought the pellet directly to the mouth) and unsuccessful trials. For each cortical location, we identified the time of peak firing rate, as well as time periods of significantly higher spike rates (>10 times SEM of randomly aligned data). The total number of identified neural units, the interquartile range (IQR), and the time period of significantly higher spike rate for each analysis are shown in Table 1.

#### Spike timing in M1 of healthy control rats

To verify that spike timing was related to the grasp event, and to compare successful vs. unsuccessful trials, we first analyzed spike activity within M1 of both hemispheres in healthy (uninjured) control rats (Group M1-C). Recordings were made during the reach task sessions on Days 3-7, 14, 21, and 28 post-implant.

First, in the M1 contralateral to the trained forelimb (i.e., contralateral M1), the peak firing rate occurred at comparable times relative to the grasp for both successful and unsuccessful trials [successful = 57 ms prior to grasp onset, Figure 3A; unsuccessful = 88 ms prior to grasp onset, Figure 3B]. Neurons in contralateral M1 displayed a comparable maximum spike rate in successful (22.44 +/- 12.88 (SEM) spikes/sec) and unsuccessful (23.85 +/- 13.24 (SEM) spikes/sec) reaches (Figure 3A). Second, during successful reaches, the criterion for significantly higher spike rate was met during the period beginning 69 ms prior to grasp onset and ending at 40 ms prior to grasp onset (Figure 3A, light blue vertical bar). This result indicates that in this preparation, contralateral M1 activity is modulated in time relative to the grasp event, as expected, and as reported by others (Hermer-Vazquez et al., 2004; Li et al., 2017). In unsuccessful reaches (Figure 3B), the average spike rate in the contralateral M1 did not exceed the threshold criterion for significantly higher spike rate at any time point during the 650 ms of the reach and grasp behavior.

In the M1 ipsilateral to the trained forelimb (i.e., ipsilateral M1), the peak firing rate occurred 130 ms *after* the grasp onset, reaching a peak spike rate of 23.3 +/- 18.4 (SEM). Significantly higher spike rates occurred during the period from 25 ms to 120 ms after grasp onset. Spike activity for ipsilateral M1 on unsuccessful trials (Figure 3D) did not exceed the criterion for significantly higher spike rates at any time during the duration of the trial.

It is important to note the distinct differences in the patterns of spiking activity between the contralateral M1 (Figure 3A) and ipsilateral M1 (Figure 3C) during successful trials in the M1-C group. The peak activity in contralateral M1 occurred 57 ms prior to the grasp onset, whereas ipsilateral M1 spiking activity reached peak activity at 130 ms *after* the grasp onset.

#### Spike timing after M1 infarct: overview of areas examined

In the PM-L and PM-C groups, neural activity was recorded for one month following ischemic injury as the rats performed the skilled reach task. In the PM-C group, neural activity was recorded in PM contralateral to the trained forelimb (i.e., contralateral PM) and in M1 ipsilateral to the trained forelimb (i.e., ipsilateral M1). In the PM-L group, neural activity was recorded in the spared PM on the side of the M1 infarct (contralateral PM/ipsilesional PM, Figure1B; shown in green) and the spared M1 in the hemisphere opposite the infarct (ipsilateral M1/contralesional M1, Figure 1B; shown in orange).

#### Spike timing after M1 infarct: contralateral (ipsilesional) PM

In the PM-C group, contralateral PM spiking profiles were similar for both successful and unsuccessful trials, in that the identified neural units were active for the duration of the motor task (successful: peak spike rate = 31.8 +/-15.6 (SEM) spikes/sec, Figure 4A; unsuccessful: peak spike rate = 29.7 +/- 14.6 (SEM) spikes/sec, Figure 4B). The criterion for significantly higher spike rate was not met at any time point during the motor task in either successful or unsuccessful trials. However, the peak spike rate occurred at 79 ms after grasp onset in successful trials and 243 ms before grasp onset in unsuccessful trials.

In contrast, in the PM-L group, spiking activity in the PM contralateral to the trained forelimb (ipsilesional PM) was not only substantially different from PM-C, but differed markedly between successful and unsuccessful trials (Figures 4C,D). For PM-L successful trials, there was a significant increase in cortical spiking activity beginning 35 ms prior to grasp onset that continued through 120 ms after grasp onset with identified neural units active (IQR) from 157 ms prior to grasp onset to 115 ms after grasp onset (Figure 4C). The peak spike rate (38.8 spikes/sec) occurred at 108 ms after grasp onset. Although spike activity for the PM-L group during unsuccessful reaches (Figure 4D) was higher than random activity for the entirety of the reach, there was no modulation or peak in activity relative to the reach task. The spike profile for PM-L unsuccessful trials exhibited a similar profile to successful and unsuccessful trials of the PM-C group.

#### Spike timing after MI infarct: ipsilateral (contralesional) M1

As expected, in the two control groups with activity recorded in M1 ipsilateral to the reaching forelimb (M1-C, PM-C), the spike trajectories on successful trials were similar. In the PM-C group, neural activity increased relative to the randomly aligned rate from 24 ms to 120 ms after grasp onset, strikingly similar to the 25 to 120 ms after grasp onset in the M1-C group (Compare Figure 3C to 5A), despite some differences in the ensemble profile.

Both PM-C and PM-L groups displayed a peak in cortical activity after grasping the pellet (successful trials). The PM-C group peak occurred at 108 ms after grasp onset (Figure 5A), averaging 39.1 +/- 21.7 (SEM) spikes/sec, while the PM-L group peak occurred at 84 ms after grasp onset (Figure 5C), averaging 30.6 +/- 24.9 (SEM) spikes/sec. After the lesion, the time of significantly increased spike activity occurred substantially earlier, beginning 122 ms *prior to* grasp onset and ending 120 ms after grasp onset (Figure 5C).

Similar to the ipsilateral M1 unsuccessful trials spike profile described for M1-C (Figure 3D), in both PM-C and PM-L groups, spike activity did not increase substantially relative to randomly aligned profiles throughout the reach trial (Figures 5B,D). Additionally, the identified neural units for PM-C and PM-L groups (PM-C = 57 units, PM-L = 107 units; Table 1) were active for comparable time intervals (PM-C IQR = 350 ms prior to grasp onset to 102 ms after grasp onset, Table 1; PM-L IQR = 322 ms prior to grasp onset to 88 ms after grasp onset, Table 1).

## DISCUSSION

The purpose of this study was to gain insight into the functional role that spared motor areas play in motor recovery. The results demonstrated that following injury to M1, the patterns of neural activity in PM contralateral to the reaching forelimb (i.e., ipsilesional PM) and in M1 ipsilateral to the reaching forelimb (i.e., contralesional M1) were altered. In contrast to the PM-C group, in which spike rates were similar throughout the reach and grasp phase of the task, the PM-L group demonstrated significantly higher spike rates beginning just before grasp onset and continuing for more than 100 ms after the grasp (Figures 4A,C). There was also a broadening of the time period over which neurons in ipsilateral M1 (i.e., contralesional M1) fired in relation to the task, with significant increases in spike rates occurring more than 100 ms before grasp onset (Figures 5A,C). To our knowledge, this is the first demonstration of a compensatory neurophysiological mechanism at the level of individual spike profiles that may explain how remote, spared cortical areas contribute to functional recovery after M1 injury.

In contralateral M1, significant increases in spike rate above randomly aligned data were seen only when data were restricted to successful trials. Since spike rate ensemble averages often appeared similar in successful and unsuccessful trials (e.g., Figures 3A,B; contralateral M1), it is possible that increasing the dataset for unsuccessful trials would have yielded a significant result. It is also possible that the lack of temporal coupling in unsuccessful trials simply reflects increased variability in the movement kinematics that occurs in unsuccessful trials, primarily during the ballistic phase of the reach. This difference may be a statistical anomaly rather than a qualitative difference in rate coding for successful vs. unsuccessful trials. Contralateral PM’s most prominent peak of spike activity occurs prior to reach onset (Hyland, 1998), supporting the notion that PM encodes some type of preparatory motor signal. In the present study, no significant elevations above randomly aligned spike rates were observed in contralateral PM. The present behavioral paradigm may have emphasized ballistic phases of forelimb movement, and thus, the timing of preparatory and planning mechanisms may have been obscured.

We observed significantly elevated spike rates in the ipsilateral M1 of control animals only after the grasp onset. Despite some differences in the appearance of the ensemble spike profiles, the time period of increased spike rate was remarkably similar in the two control conditions (M1-C: 25 to 120 ms after grasp onset; PM-C: 24 – 120 ms after grasp onset). The most parsimonious explanation for this late increase is that as the rat grasps the pellet, the non-grasping limb is actively used a) for postural control, and b) in assisting the grasping forelimb in transporting the pellet to the mouth. While these instances were not specifically quantified in the present study, postural and assistive mechanisms were clearly evident on many of the successful trials.

Following M1 lesion, significantly higher spike rates were observed in contralateral PM from 35 ms prior to grasp onset until 120 ms after grasp onset. While some modulation of spike activity prior to grasp onset might be expected (Hyland, 1998), the earlier onset of spike modulation seen only after the lesion may indicate that successful pellet retrieval required greater participation of PM in the reach phase. The substantially higher spike rates evident after the grasp onset (i.e., during the grasp phase) appear to be unique to rats with M1 lesions. These results are consistent with the notion that the PM ipsilateral to the lesion compensates for the loss of M1 and its role in movement execution. PM normally plays a critical role in prehension and grasping (Davare et al., 2008; Kantak et al., 2012). After the lesion, it is possible that a greater proportion of PM neurons are involved with the grasp phase than in control animals, and that successful retrievals are at least in part, due to a vicarious function of PM.

Changes were also evident in the M1 ipsilateral to the grasping forelimb. We observed a broadening of the time period over which significantly higher spike rates occurred. Subsequent to an M1 injury, spike activity in ipsilateral M1 increased at a substantially earlier time point (control: 24 ms *after* grasp onset; lesion: 122 ms *before* grasp onset, Figures 5A,C). It is possible that compensatory postural adjustments were made by rats after the lesion, so that the ipsilateral forelimb was used for support of the non-grasping limb, or the ipsilateral M1 may have altered its firing patterns to accommodate an alternative mechanism for movement preparation or planning.

Reorganization of cortical motor areas occurs during motor learning, but also after cortical injury. The mechanisms by which this reorganization manifests itself in the patterns of task-related neural activity are still largely unknown. Motor recovery following injury is thought to be associated with several neurophysiological (Frost et al., 2003; Shanina et al., 2006; Werhahn et al., 2003) and/or neuroanatomical (Dancause et al., 2005) changes that support restoration of function or mechanisms for motor compensation (Metz et al., 2005; Whishaw et al., 1991).

PM and M1 neural activity have overlapping characteristics with regard to spike timing, amplitude, and direction preference relative to movement during simple lever press (Saiki et al., 2014) and more complex reach and grasp (Hyland, 1998) tasks. M1 neural activity and EMG activity are strongly modulated by induced PM activity (Deffeyes et al., 2015). PM and M1 have similar efferent projections to subcortical regions (Rouiller et al., 1993). There is a high likelihood that PM is involved in the restoration of motor function following M1 injury. However, the present data suggest that it is not a simple substitution of M1 functions by PM neurons occurring during recovery. PM spike activity occurs earlier after M1 injury, consistent with a vicarious function, but is most active during the grasp phase, a result not seen in M1 of healthy animals. Thus, based on these data, PM does not recapitulate M1 function, but compensates in unique ways to accommodate recovered behavioral function.

It is likely that the alteration in contralateral PM activity associated with the reach task at least partially reflects changes in the underlying connectivity of contralateral PM and S1. Following damage to M1, axonal sprouting occurs within peri-infarct regions. In non-human primates, substantial reciprocal connections between PMv and S1 (area 1 and 2) form *de novo* after M1 injury (Dancause et al., 2005; Dancause et al., 2006). In addition, fMRI studies have demonstrated that reorganization of S1 following damage to M1 correlates with behavioral recovery (Pineiro et al., 2001), and in some cases, the reorganization suggested the function of injured M1 was shifted to S1 (Jang et al., 2002). The highest spike rates in PM after injury were observed during the grasp phase, potentially corresponding to the time when somatosensory feedback reaches S1. If new connections between S1 and PM are being formed during the recovery period, the final post-injury sensorimotor network might be shaped during such behavioral tasks. Prior *in vitro* studies have shown that growth cone development is guided by electrical activity (Ming et al., 2001). Likewise, Carmichael (2002) demonstrated that axonal sprouting generated by cortical lesions produced low-frequency synchronous neural activity patterns that were essential for initiating and driving the direction of neural sprouting. If we assume that S1 receives feedback from cutaneous and proprioceptive afferents during the grasping of the pellet at the same time period that PM reaches the peak firing rate, then the synchronous activity could potentially drive novel cortical sprouting between PM and S1. Thus, one potential neuroanatomical substrate for post-injury changes in PM during grasp may be related to greater sensorimotor integration between S1 and PM.

While changes in contralateral PM likely contributes to motor recovery directly, it is possible that the ipsilateral hemisphere also has a significant role in driving or modulating impaired motor function. This might occur through interhemispheric connections from ipsilateral M1 acting on contralateral PM or through ipsilateral control of movement. However, a secondary lesion in ipsilateral M1 to the reaching forelimb at different time points after initial infarct did not reinstate the forelimb motor deficits, suggesting that the ipsilateral M1 has a minor role in motor recovery (Gharbawie et al., 2007; Shanina et al., 2006). Other research suggests the size of ischemic infarct in M1 determines what role the ipsilateral motor cortex to reaching forelimb plays in functional recovery, with small infarcts relying on contralateral spared motor areas and large infarcts relying on ipsilateral spared motor areas to aid with behavioral recovery (Biernaskie et al., 2005).

The present study demonstrates altered spike-timing relationships in spared motor areas with respect to a behavioral task following injury. Understanding how the shift in patterns of activity that correlate with restoration of motor function will shed light on the underlying mechanisms of neural reorganization after injury. The present study is an initial step in this process, as these results suggest that spike-timing properties are altered in specific phases of the task, and that this altered activity may support compensatory movement planning and execution.

## ACKNOWLEDGMENTS

The authors thank Caleb Dunham for technical assistance. This research was supported by the National Institutes of Health (NINDS R01NS30853).

## AUTHOR CONTRIBUTIONS

Conceptualization and Methodology, A.R.P., D.J.G., and R.J.N.; Performed Research, A.R.P., S.B., and D.J.G.; Software, M.D.M.; Formal Analysis, A.R.P. and M.D.M.; Writing-Original Draft and Review & Editing, A.R.P., M.D.M., S.B., R.J.N., and D.J.G.

## References

Adkins, D.L., Voorhies, A.C., and Jones, T.A. (2004). Behavioral and neuroplastic effects of focal endothelin-1 induced sensorimotor cortex lesions. Neuroscience 128, 473–486.

Barbay, S., Guggenmos, D.J., Nishibe, M., and Nudo, R.J. (2013). Motor representations in the intact hemisphere of the rat are reduced after repetitive training of the impaired forelimb. Neurorehabil Neural Repair 27, 381–384.

Benjamin, E.J., Virani, S.S., Callaway, C.W., Chang, A.R., Cheng, S., Chiuve, S.E., Cushman, M., Delling, F.N., Deo, R., de Ferranti, S.D., et al. (2018). Heart Disease and Stroke Statistics-2018 Update: A Report From the American Heart Association. Circulation.

Biernaskie, J., Szymanska, A., Windle, V., and Corbett, D. (2005). Bi-hemispheric contribution to functional motor recovery of the affected forelimb following focal ischemic brain injury in rats. Eur J Neurosci 21, 989–999.

Bury, S.D., and Jones, T.A. (2002). Unilateral sensorimotor cortex lesions in adult rats facilitate motor skill learning with the “unaffected” forelimb and training-induced dendritic structural plasticity in the motor cortex. The Journal of neuroscience: the official journal of the Society for Neuroscience 22, 8597–8606.

Carey, J.R., Kimberley, T.J., Lewis, S.M., Auerbach, E.J., Dorsey, L., Rundquist, P., and Ugurbil, K. (2002). Analysis of fMRI and finger tracking training in subjects with chronic stroke. Brain 125, 773–788.

Dancause, N., Barbay, S., Frost, S.B., Plautz, E.J., Chen, D., Zoubina, E.V., Stowe, A.M., and Nudo, R.J. (2005). Extensive cortical rewiring after brain injury. J Neurosci 25, 10167–10179.

Dancause, N., Barbay, S., Frost, S.B., Zoubina, E.V., Plautz, E.J., Mahnken, J.D., and Nudo, R.J. (2006). Effects of small ischemic lesions in the primary motor cortex on neurophysiological organization in ventral premotor cortex. J Neurophysiol 96, 3506–3511.

Davare, M., Lemon, R., and Olivier, E. (2008). Selective modulation of interactions between ventral premotor cortex and primary motor cortex during precision grasping in humans. J Physiol 586, 2735–2742.

Deffeyes, J.E., Touvykine, B., Quessy, S., and Dancause, N. (2015). Interactions between rostral and caudal cortical motor areas in the rat. J Neurophysiol 113, 3893–3904.

Donoghue, J.P., and Wise, S.P. (1982). The motor cortex of the rat: cytoarchitecture and microstimulation mapping. J Comp Neurol 212, 76–88.

Frost, S.B., Barbay, S., Friel, K.M., Plautz, E.J., and Nudo, R.J. (2003). Reorganization of remote cortical regions after ischemic brain injury: a potential substrate for stroke recovery. J Neurophysiol 89, 32053214.

Gharbawie, O.A., Karl, J.M., and Whishaw, I.Q. (2007). Recovery of skilled reaching following motor cortex stroke: do residual corticofugal fibers mediate compensatory recovery? Eur J Neurosci 26, 3309–3327.

Grün, S., and Rotter, S. (2010). Analysis of parallel spike trains (New York: Springer).

Hermer-Vazquez, L., Hermer-Vazquez, R., Moxon, K.A., Kuo, K.H., Viau, V., Zhan, Y., and Chapin, J.K. (2004). Distinct temporal activity patterns in the rat M1 and red nucleus during skilled versus unskilled limb movement. Behav Brain Res 150, 93–107.

Hsu, J.E., and Jones, T.A. (2006). Contralesional neural plasticity and functional changes in the less-affected forelimb after large and small cortical infarcts in rats. Exp Neurol 201, 479–494.

Hyland, B. (1998). Neural activity related to reaching and grasping in rostral and caudal regions of rat motor cortex. Behav Brain Res 94, 255–269.

Jang, S.H., Han, B.S., Chang, Y., Byun, W.M., Lee, J., and Ahn, S.H. (2002). Functional MRI evidence for motor cortex reorganization adjacent to a lesion in a primary motor cortex. Am J Phys Med Rehabil 81, 844–847.

Kantak, S.S., Stinear, J.W., Buch, E.R., and Cohen, L.G. (2012). Rewiring the brain: potential role of the premotor cortex in motor control, learning, and recovery of function following brain injury. Neurorehabil Neural Repair 26, 282–292.

Kleim, J.A., Barbay, S., Cooper, N.R., Hogg, T.M., Reidel, C.N., Remple, M.S., and Nudo, R.J. (2002). Motor learning-dependent synaptogenesis is localized to functionally reorganized motor cortex. Neurobiol Learn Mem 77, 63–77.

Kleim, J.A., Barbay, S., and Nudo, R.J. (1998). Functional reorganization of the rat motor cortex following motor skill learning. J Neurophysiol 80, 3321–3325.

Li, Q., Ko, H., Qian, Z.M., Yan, L.Y.C., Chan, D.C.W., Arbuthnott, G., Ke, Y., and Yung, W.H. (2017). Refinement of learned skilled movement representation in motor cortex deep output layer. Nat Commun 8, 15834.

Liepert, J., Miltner, W.H., Bauder, H., Sommer, M., Dettmers, C., Taub, E., and Weiller, C. (1998). Motor cortex plasticity during constraint-induced movement therapy in stroke patients. Neurosci Lett 250, 5–8.

Liu, Y., and Rouiller, E.M. (1999). Mechanisms of recovery of dexterity following unilateral lesion of the sensorimotor cortex in adult monkeys. Exp Brain Res 128, 149–159.

Ludwig, K.A., Miriani, R.M., Langhals, N.B., Joseph, M.D., Anderson, D.J., and Kipke, D.R. (2009). Using a common average reference to improve cortical neuron recordings from microelectrode arrays. J Neurophysiol 101, 1679–1689.

Maccione, A., Gandolfo, M., Massobrio, P., Novellino, A., Martinoia, S., and Chiappalone, M. (2009). A novel algorithm for precise identification of spikes in extracellularly recorded neuronal signals. J Neurosci Methods 177, 241–249.

Marshall, R.S., Perera, G.M., Lazar, R.M., Krakauer, J.W., Constantine, R.C., and DeLaPaz, R.L. (2000). Evolution of cortical activation during recovery from corticospinal tract infarction. Stroke 31, 656–661.

Metz, G.A., Antonow-Schlorke, I., and Witte, O.W. (2005). Motor improvements after focal cortical ischemia in adult rats are mediated by compensatory mechanisms. Behav Brain Res 162, 71–82.

Ming, G., Henley, J., Tessier-Lavigne, M., Song, H., and Poo, M. (2001). Electrical activity modulates growth cone guidance by diffusible factors. Neuron 29, 441–452.

Miyai, I., Suzuki, T., Kang, J., Kubota, K., and Volpe, B.T. (1999). Middle cerebral artery stroke that includes the premotor cortex reduces mobility outcome. Stroke 30, 1380–1383.

Neafsey, E.J., Bold, E.L., Haas, G., Hurley-Gius, K.M., Quirk, G., Sievert, C.F., and Terreberry, R.R. (1986). The organization of the rat motor cortex: a microstimulation mapping study. Brain Res 396, 77–96.

Nhan, H., Barquist, K., Bell, K., Esselman, P., Odderson, I.R., and Cramer, S.C. (2004). Brain function early after stroke in relation to subsequent recovery. J Cereb Blood Flow Metab 24, 756–763.

Nishibe, M., Barbay, S., Guggenmos, D., and Nudo, R.J. (2010). Reorganization of motor cortex after controlled cortical impact in rats and implications for functional recovery. J Neurotrauma 27, 2221–2232.

Nudo, R.J., and Masterton, R.B. (1988). Descending pathways to the spinal cord: a comparative study of 22 mammals. J Comp Neurol 277, 53–79.

Nudo, R.J., Milliken, G.W., Jenkins, W.M., and Merzenich, M.M. (1996). Use-dependent alterations of movement representations in primary motor cortex of adult squirrel monkeys. J Neurosci 16, 785–807.

Pineiro, R., Pendlebury, S., Johansen-Berg, H., and Matthews, P.M. (2001). Functional MRI detects posterior shifts in primary sensorimotor cortex activation after stroke: evidence of local adaptive reorganization? Stroke 32, 1134–1139.

Quiroga, R.Q., Nadasdy, Z., and Ben-Shaul, Y. (2004). Unsupervised spike detection and sorting with wavelets and superparamagnetic clustering. Neural Comput 16, 1661–1687.

Rouiller, E.M., Moret, V., and Liang, F. (1993). Comparison of the connectional properties of the two forelimb areas of the rat sensorimotor cortex: support for the presence of a premotor or supplementary motor cortical area. Somatosens Mot Res 10, 269–289.

Rouiller, E.M., Yu, X.H., Moret, V., Tempini, A., Wiesendanger, M., and Liang, F. (1998). Dexterity in adult monkeys following early lesion of the motor cortical hand area: the role of cortex adjacent to the lesion. Eur J Neurosci 10, 729–740.

Saiki, A., Kimura, R., Samura, T., Fujiwara-Tsukamoto, Y., Sakai, Y., and Isomura, Y. (2014). Different modulation of common motor information in rat primary and secondary motor cortices. PLoS One 9, e98662.

Shanina, E.V., Schallert, T., Witte, O.W., and Redecker, C. (2006). Behavioral recovery from unilateral photothrombotic infarcts of the forelimb sensorimotor cortex in rats: role of the contralateral cortex. Neuroscience 139, 1495–1506.

Shimazaki, H., and Shinomoto, S. (2010a). Kernel bandwidth optimization in spike rate estimation. J Comput Neurosci 29, 171–182.

Shimazaki, H., and Shinomoto, S. (2010b). Kernel bandwidth optimization in spike rate estimation. Journal of Computational Neuroscience 29, 171–182.

Werhahn, K.J., Conforto, A.B., Kadom, N., Hallett, M., and Cohen, L.G. (2003). Contribution of the ipsilateral motor cortex to recovery after chronic stroke. Ann Neurol 54, 464–472.

Whishaw, I.Q. (1996). An endpoint, descriptive, and kinematic comparison of skilled reaching in mice (Mus musculus) with rats (Rattus norvegicus). Behav Brain Res 78, 101–111.

Whishaw, I.Q., Pellis, S.M., Gorny, B.P., and Pellis, V.C. (1991). The impairments in reaching and the movements of compensation in rats with motor cortex lesions: an endpoint, videorecording, and movement notation analysis. Behav Brain Res 42, 77–91.

